# Continuity and admixture in the last five millennia of Levantine history from ancient Canaanite and present-day Lebanese genome sequences

**DOI:** 10.1101/142448

**Authors:** Marc Haber, Claude Doumet-Serhal, Christiana Scheib, Yali Xue, Petr Danecek, Massimo Mezzavilla, Sonia Youhanna, Rui Martiniano, Javier Prado-Martinez, Michal Szpak, Elizabeth Matisoo-Smith, Holger Schutkowski, Richard Mikulski, Pierre Zalloua, Toomas Kivisild, Chris Tyler-Smith

**Author notes:** These authors contributed equally to this work (Marc Haber, Claude Doumet-Serhal, Christiana Scheib). Correspondence (M.H.), (C.T.-S.).

## Abstract

The Canaanites inhabited the Levant region during the Bronze Age and established a culture which became influential in the Near East and beyond. However, the Canaanites, unlike most other ancient Near Easterners of this period, left few surviving textual records and thus their origin and relationship to ancient and present-day populations remain unclear. In this study, we sequenced five whole-genomes from ~3,700-year-old individuals from the city of Sidon, a major Canaanite city-state on the Eastern Mediterranean coast. We also sequenced the genomes of 99 individuals from present-day Lebanon to catalogue modern Levantine genetic diversity. We find that a Bronze Age Canaanite-related ancestry was widespread in the region, shared among urban populations inhabiting the coast (Sidon) and inland populations (Jordan) who likely lived in farming societies or were pastoral nomads. This Canaanite-related ancestry derived from mixture between local Neolithic populations and eastern migrants genetically related to Chalcolithic Iranians. We estimate, using linkage-disequilibrium decay patterns, that admixture occurred 6,600-3,550 years ago, coinciding with massive population movements in the mid-Holocene triggered by aridification ~4,200 years ago. We show that present-day Lebanese derive most of their ancestry from a Canaanite-related population, which therefore implies substantial genetic continuity in the Levant since at least the Bronze Age. In addition, we find Eurasian ancestry in the Lebanese not present in Bronze Age or earlier Levantines. We estimate this Eurasian ancestry arrived in the Levant around 3,750-2,170 years ago during a period of successive conquests by distant populations such as the Persians and Macedonians.

---

The Near East, including the Levant, has been central to human prehistory and history from the expansion out of Africa 50-60 thousand years ago (kya),^1^ through post-glacial expansions^2^ and the Neolithic transition 10 kya, to the historical period when Ancient Egyptians, Greeks, Phoenicians, Assyrians, Babylonians, Persians, Romans and many others left their impact on the region.^3^ Aspects of the genetic history of the Levant have been inferred from present-day DNA,^4; 5^ but the more comprehensive analyses performed in Europe^6-11^ have shown the limitations of relying on present-day information alone, and highlighted the power of ancient DNA (aDNA) for addressing questions about population histories.^12^ Unfortunately, although the few aDNA results from the Levant available so far are sufficient to reveal how much its history differs from that of Europe,^13^ more work is needed to establish a thorough understanding of Levantine genetic history. Such work is hindered by the hot and sometimes wet environment,^12; 13^ but improved aDNA technologies including use of the petrous bone as a source of DNA^14^ and the rich archaeological remains available, encouraged us to further explore the potential of aDNA in this region. Here, we present genome sequences from five Bronze Age Lebanese samples and show how they improve our understanding of the Levant’s history over the last five millennia.

During the Bronze Age in the Levant, around 3-4 kya, a distinctive culture emerged as a Semitic-speaking people known as the Canaanites. The Canaanites inhabited an area bounded by Anatolia to the north, Mesopotamia to the East, and Egypt to the south, with access to Cyprus and the Aegean through the Mediterranean. Thus the Canaanites were at the centre of emerging Bronze Age civilizations and became politically and culturally influential.^15^ They were later known to the ancient Greeks as the Phoenicians who, 2.3-3.5 kya, colonized territories throughout the Mediterranean reaching as far as the Iberian Peninsula.^16^ However, for uncertain reasons, but perhaps related to the use of papyrus instead of clay for documentation, few textual records have survived from the Canaanites themselves and most of their history known today has been reconstructed from ancient Egyptian and Greek records, the Hebrew Bible and archaeological excavations.^15^ Many uncertainties still surround the origin of the Canaanites: Ancient Greek historians believed their homeland was located in the region of the Persian Gulf.^16; 17^ However, modern researchers tend to reject this hypothesis because of archaeological and historical evidence of population continuity through successive millennia in the Levant. The Canaanite culture is alternatively thought to have developed from local Chalcolithic people who were themselves derived from people who settled in farming villages in the 9-10 kya during the Neolithic period.^15^ Uncertainties also surround the fate of the Canaanites: the Bible reports the destruction of the Canaanite cities and the annihilation of its people; if true, the Canaanites could not have directly contributed genetically to present-day populations. However, no archaeological evidence has so far been found to support widespread destruction of Canaanite cities between the Bronze and Iron Ages: cities on the Levant coast such as Sidon and Tyre show continuity of occupation until the present day.

aDNA research has the potential to resolve many questions related to the history of the Canaanites, including their place of origin and fate. Here, we sampled the petrous portion of temporal bones belonging to five ancient individuals dated to between 3,750 and 3,650 years ago (ya) from Sidon, which was a major Canaanite city-state during this period (Figure S1 and S2). We extracted DNA and built double-stranded libraries according to published protocols.^14 18-20^ We sequenced the libraries on an Illumina HiSeq 2500 using 2×75 bp reads and processed the sequences using the PALEOMIX pipeline.^21^ We retained reads ≥ 30bp and collapsed pairs with minimum overlap of 15bp, allowing a mismatch rate of 0.06 between the pairs. We mapped the merged sequences to the ***hs37d5*** reference sequence, removed duplicates, removed two bases from the ends of each read, and randomly sampled a single sequence with a minimum quality of ≥20 to represent each SNP. We obtained a genomic coverage of 0.4-2.3x and a mitochondrial DNA (mtDNA) genome coverage of 53-164x (Table 1). In order to assess ancient DNA authenticity, we estimated X-chromosome contamination^22; 23^(Table S1) and restricted some analyses to sequences with aDNA damage patterns^24; 25^ (Figure S3 and S4), and as a result demonstrate that the sequence data we present are endogenous and minimally contaminated.

**Table 1:**
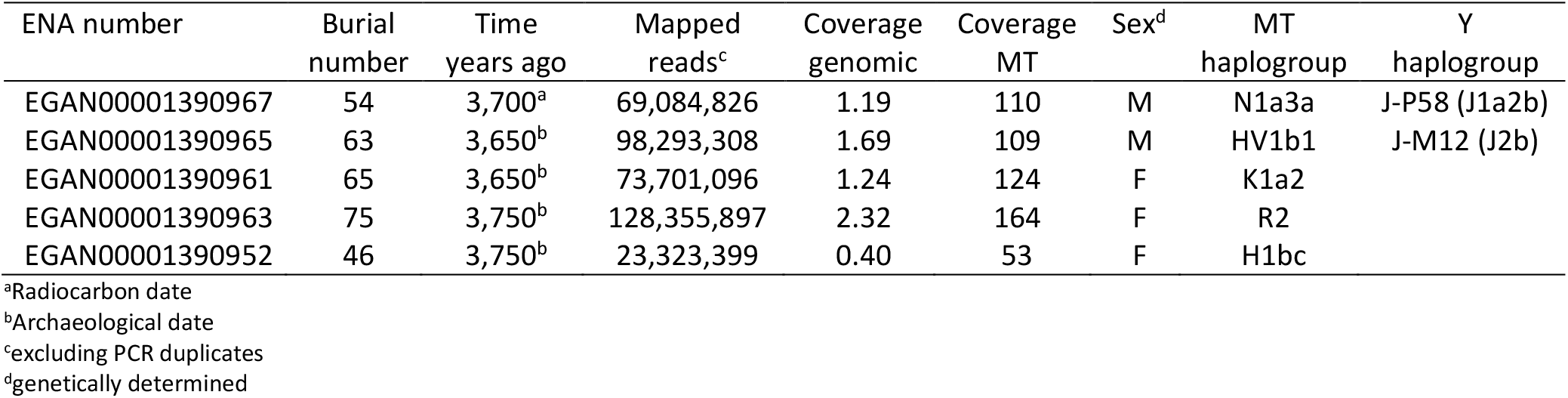
Samples analysed in this study

| ENA number | Burial number | Time years ago | Mapped reads<sup>c</sup> | Coverage genomic | Coverage MT | Sex<sup>d</sup> | MT haplogroup | Y haplogroup |
| --- | --- | --- | --- | --- | --- | --- | --- | --- |
| EGAN00001390967 | 54 | 3,700<sup>a</sup> | 69,084,826 | 1.19 | 110 | M | N1a3a | J-P58 (J1a2b) |
| EGAN00001390965 | 63 | 3,650<sup>b</sup> | 98,293,308 | 1.69 | 109 | M | HV1b1 | J-M12 (J2b) |
| EGAN00001390961 | 65 | 3,650<sup>b</sup> | 73,701,096 | 1.24 | 124 | F | K1a2 |  |
| EGAN00001390963 | 75 | 3,750<sup>b</sup> | 128,355,897 | 2.32 | 164 | F | R2 |  |
| EGAN00001390952 | 46 | 3,750<sup>b</sup> | 23,323,399 | 0.40 | 53 | F | H1bc |  |

<sup>a</sup>Radiocarbon date<sup>b</sup>Archaeological date<sup>c</sup>excluding PCR duplicates<sup>d</sup>genetically determined

Additionally, we sequenced whole-genomes of 99 present-day Lebanese individuals to ~8x coverage on an Illumina HiSeq 2500 using 2× 100 bp reads. We merged the low-coverage Lebanese data with four high-coverage (30x) Lebanese samples,^26^ 1000 Genomes Project phase 3 CEU, YRI, and CHB populations,^27^ and sequence data previously published from regional populations (Egyptians, Ethiopians and Greeks).^1; 26^ Raw calls were generated using bcftools (bcftools mpileup -C50 -pm3 -F0.2 -d10000 | bcftools call -mv, version 1.2-239-g8749475) and filtered to include only SNPs with the minimum of 2 alternate alleles in at least one population and site quality larger than 10; we excluded sites with a minimum per-population HWE and total HWE less than 0.01^28^ and sites within 3bp of an indel. The filtered calls were then pre-phased using shapeit (v2.r790)^29^ and their genotypes refined using beagle (v4.1).^30^

We combined our ancient and modern samples with previously published ancient data^6-11; 13; 24; 31; 32^ (Figure 1A) resulting in a dataset of 389 individuals and 1,046,317 SNPs when ancient and Lebanese samples were analysed, and 546,891 SNPs when 2,583 modern samples from the Human Origins genotype data were included in the analysis.^9; 33^ The ancient samples were grouped following the labels assigned by Lazaridis et al. ^13^ on the basis of archaeological culture, chronology and genetic clustering. We used this dataset to shed light on the genetic history of the Canaanites, resolving their relationship to other ancient populations and assessing their genetic contribution to present-day populations.

**Figure 1.**
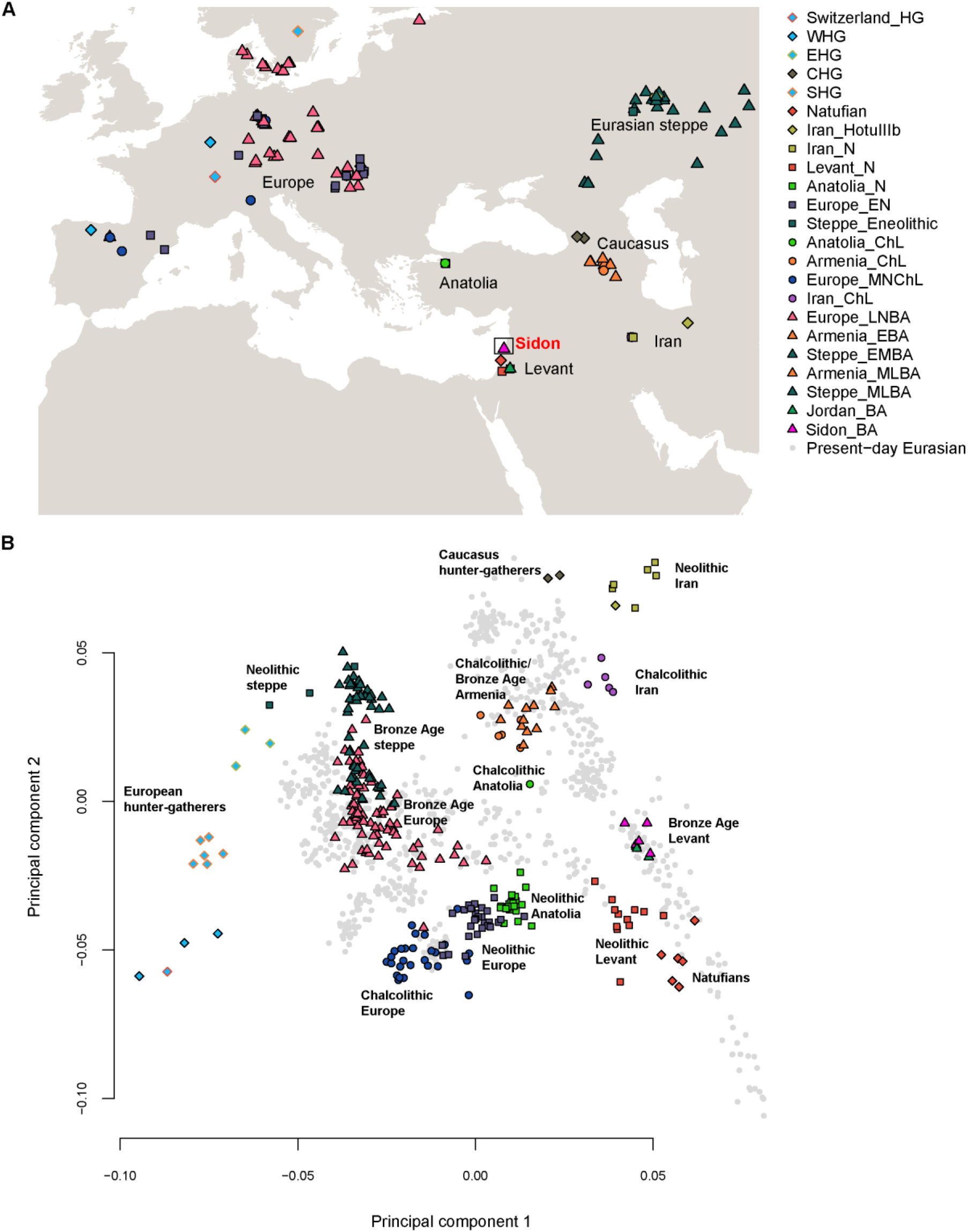
Population locations and genetic structure. (A) The map shows the location of the newly sequenced Bronze Age Sidon samples (pink triangle labelled with red text), as well as the locations of published ancient samples used as comparative data in this study. (B) PCA of ancient Eurasian samples (colored shapes) projected using eigenvectors from present-day Eurasian populations (grey points).

We first explored our dataset using principal component analysis (PCA)^34^ on present-day West Eurasian (including Levantine) populations and projected the ancient samples onto this plot (Figure 1B and S5). The Bronze Age Sidon samples (Sidon_BA) overlap with present-day Levantines and were positioned between the ancient Levantines (Natufians/Neolithic) and ancient Iranians (Neolithic/Chalcolithic). The overlap between the Bronze Age and present-day Levantines suggests a degree of genetic continuity in the region. We explored this further by computing the statistic *f*4(Lebanese, present-day Near Easterner; Sidon_BA, Chimpanzee) using *qpDstat*^33^(with parameter f4mode: YES) and found Sidon_BA shared more alleles with the Lebanese than with most other present-day Levantines (Figure S6), supporting local population continuity as observed in Sidon’s archaeological records. When we substituted present-day Near Easterners with a panel of 150 present-day populations available in the Human Origins dataset, we found only Sardinians and Italian_North shared significantly more alleles with Sidon_BA compared with the Lebanese (Figure S7). Sardinians are known to have retained a large proportion of ancestry from Early European farmers (EEF) and therefore the increased affinity to Sidon_BA could be related to a shared Neolithic ancestry. We computed *f*4(Lebanese, Sardinian/Italian_North; Sidon_BA, Levant_N) and found no evidence of increased affinity of Sardinians or Italian_North to Sidon_BA after the Neolithic (both Z-scores are positive). We next wanted to explore if the increased affinity of Sidon_BA to the Lebanese could also be observed when analysing functionally important regions of the genome which are less susceptible to genetic drift. Our sequence data allowed us to scan loci linked to phenotypic traits and loci previously identified as functional variants in the Lebanese and other Levantines.^35-37^ Using a list of 84 such variants (Table S2), we estimated the allele frequency (AF) in Sidon_BA using ANGSD^22^ based on a method from Li et al.^38^ and calculated Pearson pair-wise correlation coefficients between AF in Sidon_BA and AF in Africans, Europeans, Asians^27^ and Lebanese. We found a high significant correlation between Sidon_BA and the Lebanese (r = 0.74; 95% CI = 0.63-0.82; p value = 8.168e-16) and lower correlations between Sidon_BA and Europeans (r = 0.56), Africans, (r = 0.55) and Asians (r = 0.53) (Figure S8). These results support substantial local population continuity and suggest that several present-day genetic disorders might stem from risk alleles which were already present in the Bronze Age population. In addition, SNPs associated with phenotypic traits show Sidon_BA and the Lebanese had comparable skin, hair, and eye colours (in general: light intermediate skin pigmentation, brown eyes and dark hair) with similar frequencies of the underlying causal variants in ***SLC24A5*** and ***HERC2,*** but with Sidon_BA probably having darker skin than Lebanese today from variants in ***SLC45A2*** resulting in darker pigmentation (Table S2).

The PCA shows that Sidon_BA clusters with three individuals from Early Bronze Age Jordan (Jordan_BA) found in a cave above the Neolithic site of ‘Ain Ghazal and probably associated with an Early Bronze Age village close to the site.^13^ This suggests that people from the highly differentiated urban culture on the Levant coast and inland people with different modes of subsistence were nevertheless genetically similar, supporting previous reports that the different cultural groups who inhabited the Levant during the Bronze Age, such as the Ammonites, Moabites, Israelites and Phoenicians, each achieved their own cultural identities but all shared a common genetic and ethnic root with Canaanites.^15^ Lazaridis et al.^13^ reported that Jordan_BA can be modelled as mixture of Neolithic Levant (Levant_N) and Chalcolithic Iran (Iran_ChL). We computed the statistic *f*4(Levant_N, Sidon_BA; Ancient Eurasian, Chimpanzee) and found populations from the Caucasus and Iran shared more alleles with Sidon_BA than with Neolithic Levant (Figure 2A). We then used *qpAdm*^8^ (with parameter allsnps: YES) to test if Sidon_BA can be modelled as mixture of Levant_N and any other ancient population in the dataset and found good support for the model of Sidon_BA being a mixture of Levant_N (48.4± 4.2%) and Iran_ChL (51.6± 4.2%) (Figure 2B; Table S3).

**Figure 2.**
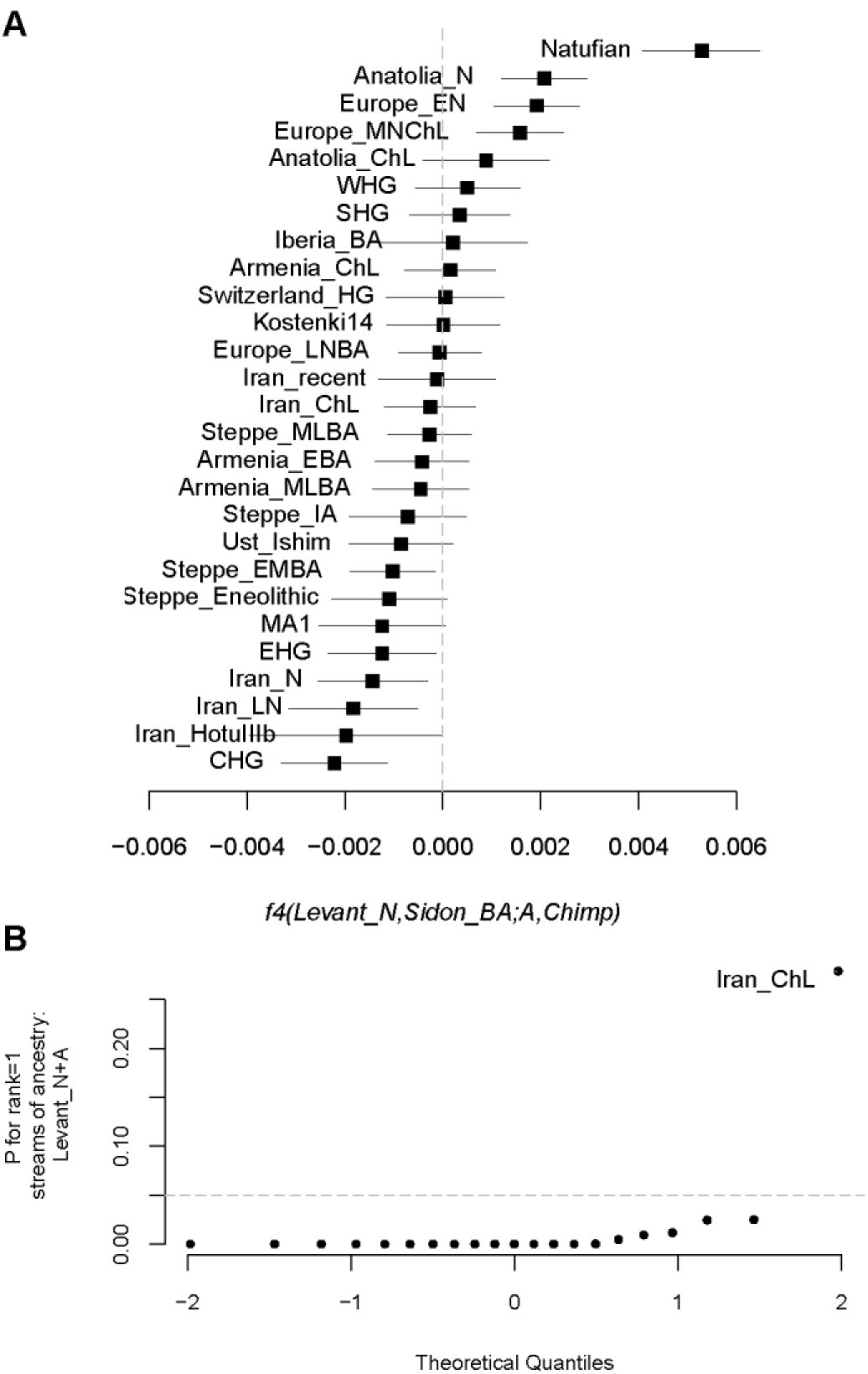
Admixture in Bronze Age Levantine populations. (A) The statistic *f*4(Levant_N, Sidon_BA; Ancient Eurasian, Chimpanzee) is most negative for populations from the Caucasus and Iran suggesting an increase in ancestry related to these populations in Sidon after the Neolithic period. (B) Modelling Sidon as mixture between Neolithic Levant and an ancient Eurasian population shows that Chalcolithic Iran fits the model best when using a large number of outgroups: Ust_Ishim, Kostenki14, MA1, Han, Papuan, Ami, Chukchi, Karitiana, Mbuti, Switzerland_HG, EHG, WHG, and CHG. Sidon_BA can then be modelled using ***qpAdm*** as 0.484± 0.042 Levant_N and 0.516± 0.042 Iran_ChL.

In addition, the two Sidon_BA males carried the Y-chromosome haplogroups^39^ J-P58 (J1a2b) and J-M12 (J2b) (Table 1 and S4; Figure S9), both common male lineages in the Near East today. We compiled frequencies of Y-chromosomal haplogroups in this geographical area and their changes over time in a dataset of ancient and modern Levantine populations (Figure S10), and note, similarly to Lazaridis et al.,^13^ that haplogroup J was absent in all Natufian and Neolithic Levant male individuals examined thus far, but emerged during the Bronze Age in Lebanon and Jordan along with ancestry related to Iran. All five Sidon_BA individuals had different mitochondrial DNA haplotypes^40^ (Table 1), belonging to paragroups common in present-day Lebanon and nearby regions (Table S5) but with additional derived variants not observed in our present-day Lebanese dataset.

We next sought to estimate the time when the Iranian ancestry penetrated the Levant. Our results support genetic continuity since the Bronze Age and thus our large dataset of present-day Lebanese provided an opportunity to explore the admixture time using admixture-induced linkage disequilibrium (LD) decay. Using ALDER^41^ (with mindis: 0.005), we set the Lebanese as the admixed test population and Natufians, Levant_N, Sidon_BA, Iran_N, and Iran_ChL as reference populations. To account for the small number of individuals in the reference populations and the limited number of SNPs in the dataset, we took a lenient minimum Z-score=2 to be suggestive of admixture. The most significant result was for mixture of Levant_N and Iran_ChL (p=0.013) around 181 ± 54 generations ago, or ~5,000 ± 1,500 ya assuming a generation time of 28 years (Figure S11A). This admixture time, based entirely on genetic data, fits the known ages of the samples based on archaeological data since it falls between the dates of Sidon_BA (3,650-3,750 ya) and Iran_ChL (6,500-5,500 ya). The admixture time also overlaps with the rise and fall of the Akkadian Empire which controlled the region from Iran to the Levant between ~4.4 and 4.2 kya. The Akkadian collapse is argued to have been the result of a widespread aridification event around 4,200 ya, possibly caused by a volcanic eruption.^42; 43^ Archaeological evidence in this period documents large-scale influxes of refugees from Northern Mesopotamia towards the south, where cities and villages became overpopulated.^44^ Future sampling of ancient DNA from Northern Syria and Iraq may reveal if these migrants carried the Iran_ChL-related ancestry we observe in Bronze Age Sidon and Jordan.

Although ***f4*** tests showed that present-day Lebanese share significantly more alleles with Sidon_BA than other Near Eastern populations do, indicating genetic continuity, we failed to model the present-day Lebanese using streams of ancestry coming only from Levant_N and Iran_ChL (*qpAdm* rank1 p= 8.36E-07), in contrast to our success with Sidon_BA. We therefore further explored our dataset by running ADMIXTURE^45^ in a supervised mode using Western hunter-gatherers (WHG), Eastern hunter-gatherers (EHG), Levant_N, and Iran_N as reference populations. These four populations have been previously^13^ found to contribute genetically to most West Eurasians. The ADMIXTURE results replicate the findings from ***qpAdm*** for Sidon_BA and show mixture of Levant_N and Iranian populations (Figure 3A). However, the present-day Lebanese, in addition to their Levant_N and Iranian ancestry, have a component (11-22%) related to EHG and Steppe populations not found in Bronze Age populations (Figure 3A). We confirm the presence of this ancestry in the Lebanese by testing *f*4(Sidon_BA, Lebanese; Ancient Eurasian, Chimpanzee) and find that Eurasian hunter-gatherers and Steppe populations share more alleles with the Lebanese than with Sidon_BA (Figure 3B). We next tested a model of the present-day Lebanese as a mixture of Sidon_BA and any other ancient Eurasian population using ***qpAdm.*** We found that the Lebanese can be best modelled as Sidon_BA 93±1.6% and a Steppe Bronze Age population 7±1.6% (Figure 3C; Table S6). To estimate the time when the Steppe ancestry penetrated the Levant we used, as above, LD-based inference and set the Lebanese as admixed test population with Natufians, Levant_N, Sidon_BA, Steppe_EMBA, and Steppe_MLBA as reference populations. We found support (p=0.00017) for a mixture between Sidon_BA and Steppe_EMBA which has occurred around 2,950±790 ya (Figure S11B). It is important to note here that Bronze Age Steppe populations used in the model need not be the actual ancestral mixing populations, and the admixture could have involved a population which was itself admixed with a Steppe-like ancestry population. The time period of this mixture overlaps with the decline of the Egyptian empire and its domination over the Levant, leading some of the coastal cities to thrive, including Sidon and Tyre, which established at this time a successful maritime trade network throughout the Mediterranean. The decline in Egypt’s power was also followed by a succession of conquests of the region by distant populations such as the Assyrians, Persians, and Macedonians, any or all of whom could have carried the Steppe-like ancestry observed here in the Levant after the Bronze Age.

**Figure 3.**
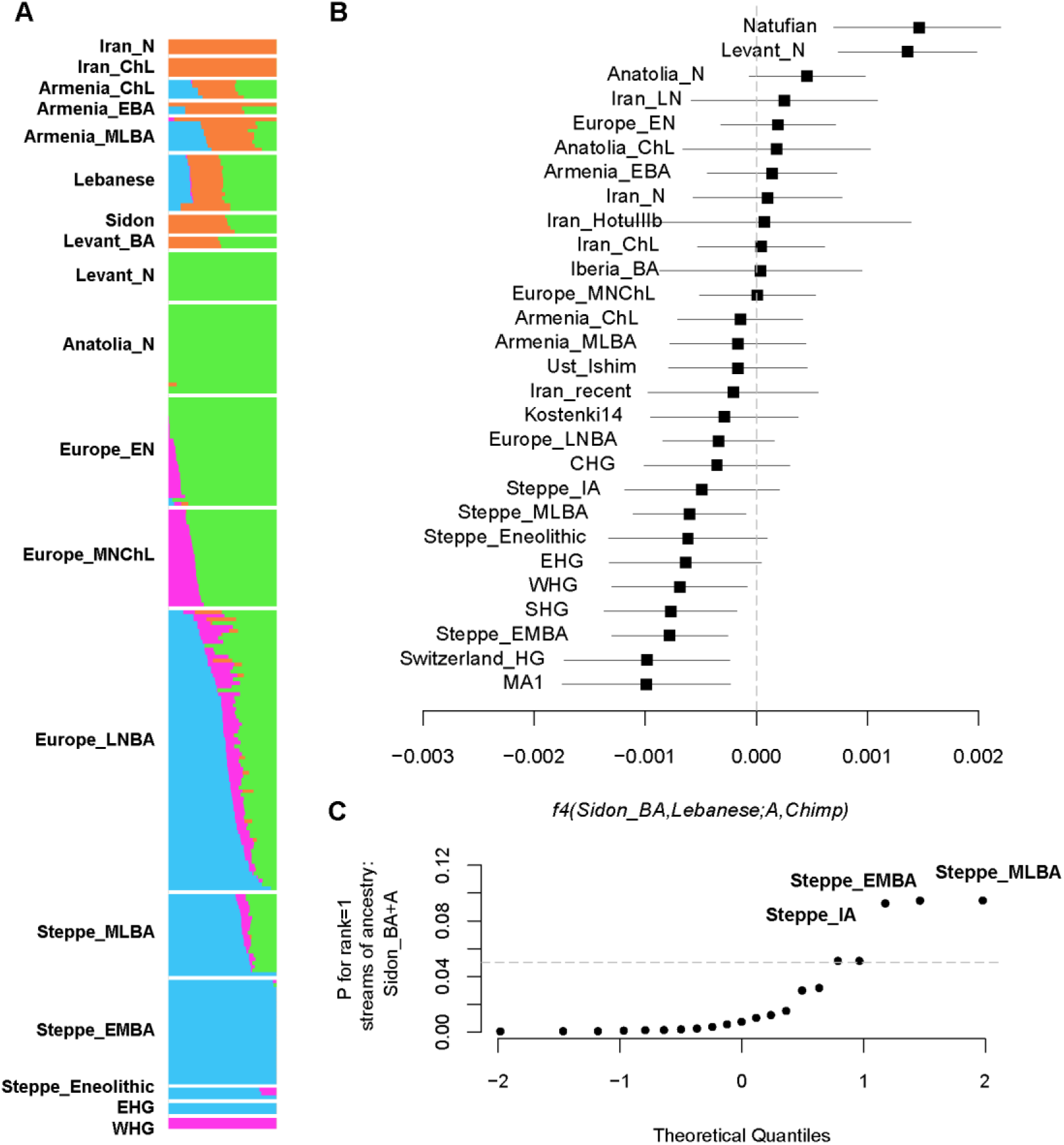
Admixture in present-day Levantine populations. (A) Supervised ADMIXTURE using Levant_N, Iran_N, EHG and WHG as populations with fixed ancestries. A Eurasian ancestry found in Eastern hunter-gatherers and the steppe Bronze Age appears in present-day Levantines after the Bronze Age. (B) The statistic *f*4(Sidon_BA, Lebanese; Ancient Eurasian, Chimpanzee) confirms the ADMIXTURE results and is most negative for populations from the steppe and Eurasian hunter-gatherers. (C) Present-day Lebanese can be modelled as mixture between Bronze Age Sidon and a steppe population. The model with mix proportions 0.932±0.016 Sidon_BA and 0.068±0.016 steppe_EMBA for Lebanese is supported with the lowest SE.

In this report we have analysed the first ancient whole-genome sequence data from a Levantine civilization, and provided insights into how the Bronze Age Canaanites were related to other ancient populations and how they have contributed genetically to present-day ones (Figure 4). Many of our inferences rely on the limited number of ancient samples available, and we are only just beginning to reconstruct a genetic history of the Levant or the Near East as thorough as that of Europeans who, in comparison, have been extensively sampled. In the future, it will be important to examine samples from the Chalcolithic/Early Bronze Age Near East to understand the events leading to admixture between local populations and the eastern migrants. It will also be important to analyze samples from the Iron Age to trace back the Steppe-like ancestry we find today in present-day Levantines. Our current results show that such studies are feasible.

**Figure 4.**
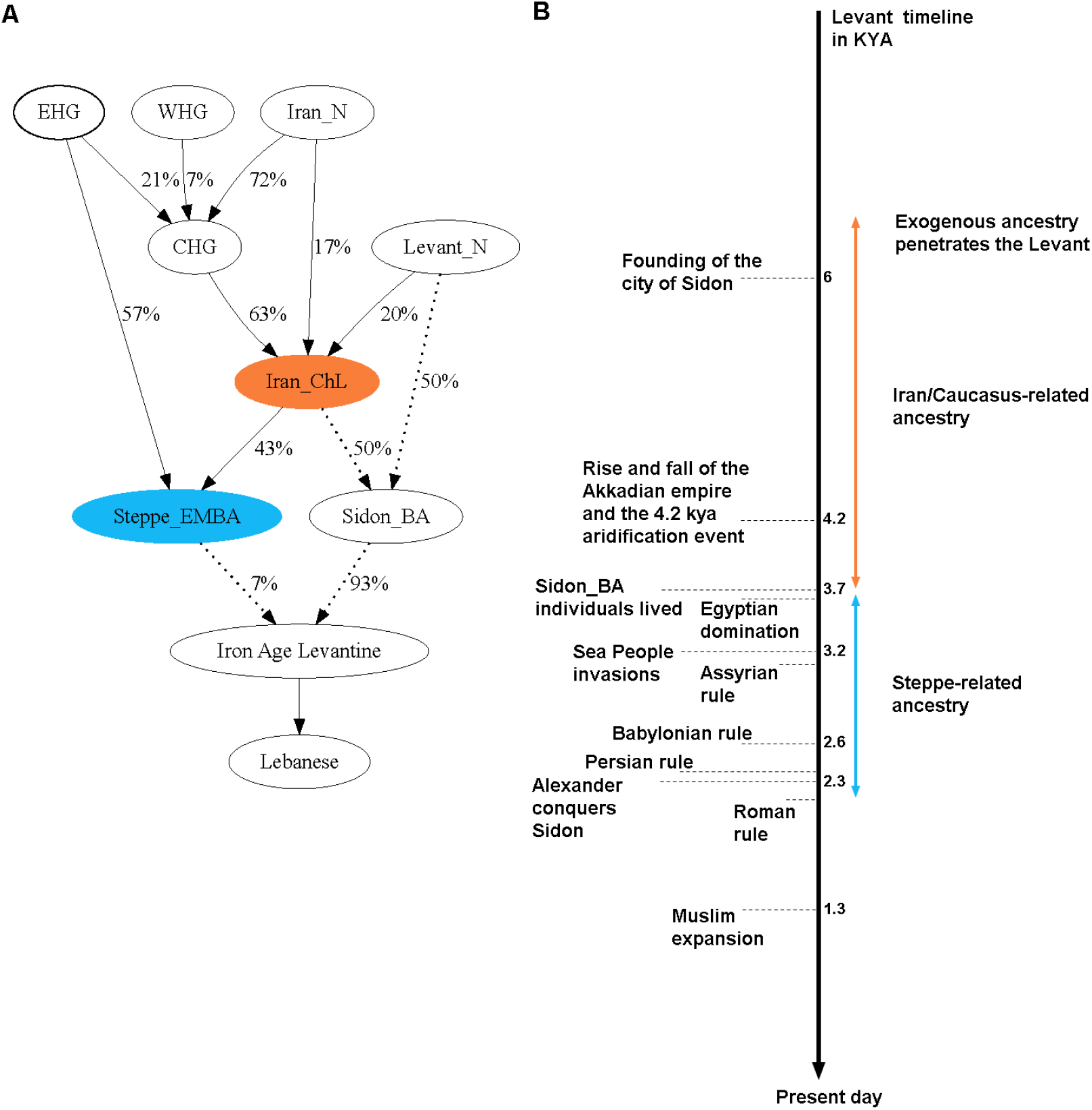
Genetic history of the Levant. (A) A model of population relationships which fits the ***qpAdm*** results from Lazaridis et al.^13^ (solid arrows) and this study (dotted arrows). Percentages on arrows are the inferred admixture proportions. (B) Levant timeline of historical events with genetically inferred admixture dates shown as coloured double-ended arrows with length representing the SE.

## Acknowledgements

We thank the present-day donors who contributed their samples to this study. M.H., Y.X., P.D., R.M., J.P.-M., M.S. and C.T.-S. were supported by The Wellcome Trust (098051).

